# Transcriptional Activation of Elephant Shark Mineralocorticoid Receptor by Corticosteroids, Progesterone and Spironolactone

**DOI:** 10.1101/265348

**Authors:** Yoshinao Katsu, Satomi Kohno, Kaori Oka, Xiaozhi Lin, Sumika Otake, Nisha E. Pillai, Wataru Takagi, Susumu Hyodo, Byrappa Venkatesh, Michael E. Baker

## Abstract

We report the analysis of activation of full-length mineralocorticoid receptor (MR) from elephant shark, a cartilaginous fish belonging to the oldest group of jawed vertebrates by corticosteroids and progesterone. Based on their measured activities, aldosterone, cortisol, 11-deoxycorticosterone, corticosterone, 11-deoxcortisol, progesterone and 19-norprogesterone are potential physiological mineralocorticoids. However, aldosterone, the physiological mineralocorticoid in humans and other terrestrial vertebrates, is not found in cartilaginous or ray-finned fishes. Although progesterone activates ray-finned fish MRs, progesterone does not activate human, amphibian or alligator MRs, suggesting that during the transition to terrestrial vertebrates, progesterone lost the ability to activate the MR. Both elephant shark MR and human MR are expressed in the brain, heart, ovary, testis and other non-epithelial tissues, indicating that MR expression in diverse tissues evolved in the common ancestor of jawed vertebrates. Our data suggest that progesterone-activated MR may have unappreciated functions in elephant shark ovary and testis.

## Introduction

The mineralocorticoid receptor (MR) belongs to the nuclear receptor family, a large and diverse group of transcription factors that also includes receptors for glucocorticoids (GR), progesterone (PR) androgens (AR) and estrogens (ER) (*1, 2*). Sequence analysis has revealed that the MR and GR are closely related (*3*); phylogenetic analysis indicates that MR and GR evolved from a corticosteroid receptor (CR) that was present in jawless vertebrates, such as lamprey and hagfish (*4-7*). A distinct mineralocorticoid receptor (MR) first appears in cartilaginous fishes (Chondrichthyes), the oldest group of extant jawed vertebrates (gnathostomes) that diverged from bony vertebrates about 450 million years. As such, cartilaginous fishes are a crucial group in understanding the origin and evolution of jawed vertebrate morphology and physiology (*8, 9*). Like mammals, cartilaginous fishes contain the full complement of adrenal and sex steroid receptors: AR, ER, GR, MR and PR (*1, 2, 4, 10*).

Aldosterone (Aldo) is the physiological activator of transcription of human MR in epithelial tissues, such as the kidney distal collecting tubules and the colon, in which the MR regulates electrolyte homeostasis (*6, 11-14*). The MR also is found in brain, heart, aorta, lung, liver, spleen, adipose tissue, testis, breast and ovary (*12–20*), tissues in which the MR is not likely to regulate electrolyte homeostasis, its classical function. The physiological function of the MR in these tissues is still being elucidated (*14, 18, 20, 21*).

The MR and other steroid receptors have a characteristic modular structure consisting of an N-terminal domain (NTD) (domains A and B), a central DNA-binding domain (DBD) (domain C), a hinge domain (D) and a C-terminal ligand-binding domain (LBD) (domain E) (*2, 4, 22–24*) (Figure 1). The LBD alone is competent to bind steroids (*4, 22, 25–27*). However, interactions between the NTD (A/B domains) and the LBD and coactivators are important regulators of transcriptional activation of mammalian MRs (*28–35*) and ray-finned fish MRs (*34–36*). Intriguingly, GAL-DBD-hinge-LBD constructs of zebrafish MR have different responses to progesterone (Prog) and some corticosteroids than do GAL-DBD-hinge-LBD constructs of human, chicken, alligator and Xenopus MRs (*35*).

**Figure 1.**
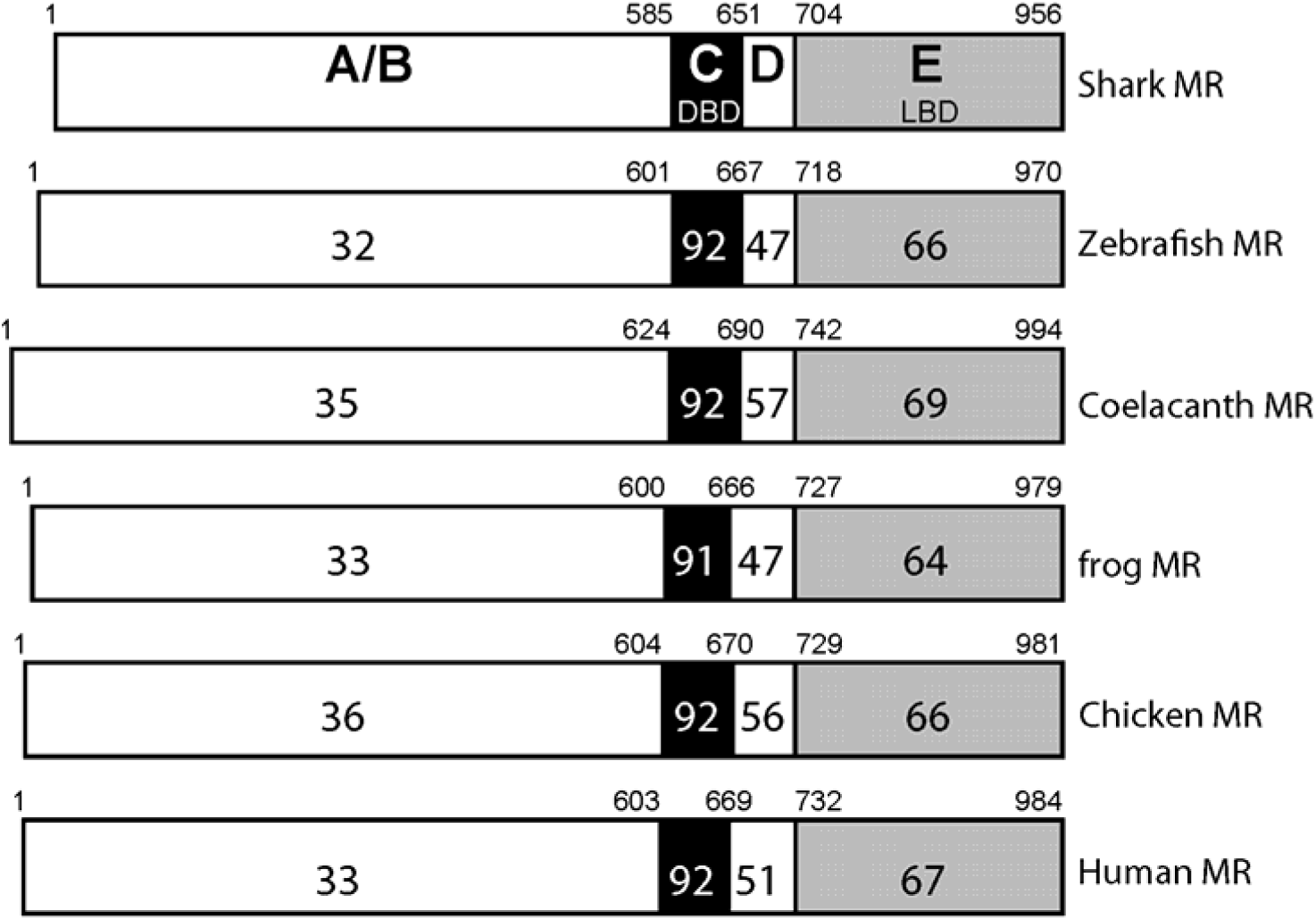
Comparison of domains in elephant shark MR with vertebrate MRs. MRs from elephant shark (shark), zebrafish, coelacanth, Xenopus (frog), chicken and human are compared. The functional A/B domain to E domains are schematically represented with the numbers of amino acid residues and the percentage of amino acid identity is depicted. GenBank accession numbers: elephant shark MR (XP_007902220), zebrafish MR (NP_001093873), coelacanth MR (XP_014348128), Xenopus (NP_001084074), chicken (ACO37437), human MR (NP_000892).

The timing of the evolution of this difference in transcriptional activation between full-length and truncated MRs in ray-finned fishes and terrestrial vertebrates, as well as when expression of the MR in non-epithelial tissues evolved, are not known. Also unresolved is the identity of the ancestral mineralocorticoid in cartilaginous fishes and the current mineralocorticoid in ray-finned fish because aldosterone (Aldo), the physiological mineralocorticoid in terrestrial vertebrates, is not found in either cartilaginous fishes or ray-finned fishes. Aldo first appears in lungfish, a lobe-finned fish (*37*), which are forerunners of terrestrial vertebrates (*38*). Thus, the identity of the physiological mineralocorticoid in cartilaginous fishes and ray-finned fishes is unknown, although cortisol and 11-deoxycorticosterone have been proposed as candidates (*39–45*).

Complicating the identity of the physiological mineralocorticoid in cartilaginous and ray-finned fishes is evidence that progesterone (Prog) and 19-norprogesterone (19norProg), along with spironolactone (Spiron) (Figure 2), are transcriptional activators of several ray-finned fish MRs (*24, 36, 44*), including zebrafish MR (*35, 46*), and of chicken MR (*35*). In contrast, these steroids are antagonists for human MR (*25, 27, 47*), alligator MR and Xenopus MR (*35*). Ray-finned fish MRs and chicken MR differ in their response to Prog, 19norProg and Spiron, raising the question of whether the response to Prog and Spiron evolved in ray-finned fish before or after the divergence of ray-finned fish from the lobe-finned fish lineage that led to tetrapods.

**Figure 2.**
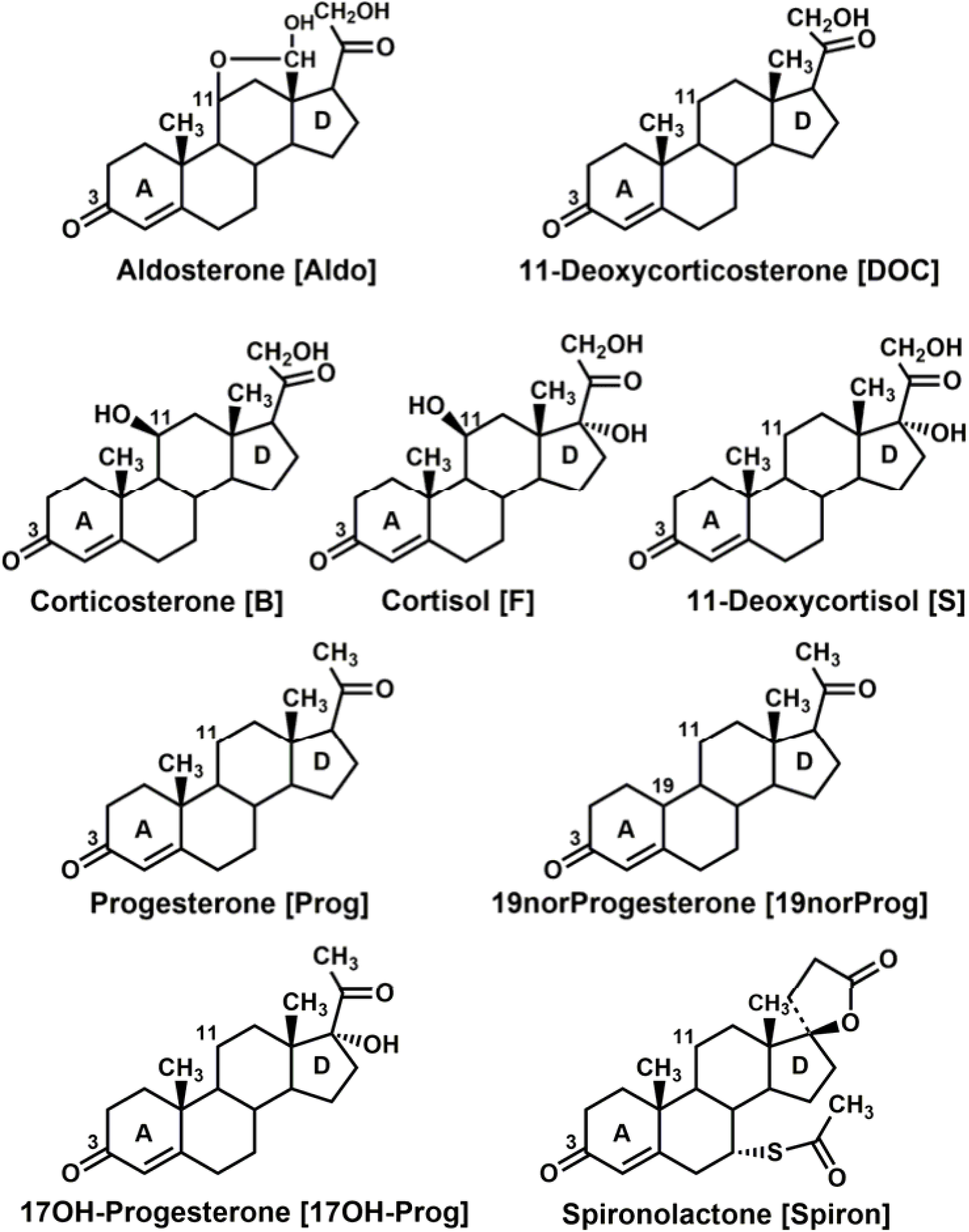
Structures of steroids that are ligands for the MR. Aldo, 11-deoxycorticosterone and S are physiological mineralocorticoids in terrestrial vertebrates (*4, 6, 12, 48*). S is both a mineralocorticoid and a glucocorticoid in lamprey (*4, 49*) and a glucocorticoid in ray-finned fish (*50*). Cortisol is a physiological glucocorticoid in terrestrial vertebrates and ray-finned fish (*4, 10, 51–53*). corticosterone is a glucocorticoid in rats and mice (*4*). Aldo, 11-deoxycorticosterone, cortisol, corticosterone and Prog have a similar high affinity for human MR (*3, 54-56*). Prog, 19norProg, 17OH-Prog and Spiron are antagonists for human MR (*24, 47, 54*) and rat MR (*57, 58*). Prog, 19norProg, and Spiron are agonists for fish MRs (*24, 36, 44*). 19norProg is a weak agonist for rat MR (*59, 60*).

Transcriptional activation by corticosteroids and other 3-ketosteroids of a full-length cartilaginous fish MR has not been investigated. Only truncated skate MR (*61*), consisting of the GAL4-DBD fused to the D domain and E domain of the MR (MR-LBD) has been studied for its response to corticosteroids. In these studies, Carroll et al. found that Aldo, corticosterone, 11-deoxycorticosterone, and cortisol (Figure 2) are transcriptional activators of truncated skate MR.

Elephant shark MR is an attractive receptor to study early events in the evolution of mechanisms for regulating MR transcription because, in addition to its phylogentic position as a sister group of bony vertebrates, genomic analyses reveal that elephant shark genes are evolving slowly (*9*), making their genes a window into the past. We therefore investigated transcriptional activation of full-length and truncated elephant shark MR by Aldo, 11-deoxycorticosterone, corticosterone, 11-deoxycortisol, cortisol, Prog, 19norProg, 17-hydroxyprogesterone (17OH-Prog) and Spiron. Interestingly, all 3-ketosteroids, including progesterone, had half-maximal responses (EC50s) of 1 nM or lower for full-length elephant shark MR. Transcriptional activation by Prog, 19norProg and Spiron of truncated elephant shark MR resembled that of zebrafish MR and not chicken MR, indicating that Prog activation of MR is an ancestral response, conserved in cartilaginous fishes and ray-finned fish, but lost in Xenopus, alligator and human MRs, and distinct from activation of chicken MR, which arose independently. We investigated expression levels of elephant shark MR by using RNA-seq data and find widespread expression of MR in various elephant shark tissues (gill, kidney, heart, intestine, liver, spleen, brain, ovary and testis). This indicates that widespread expression of human MR in tissues, such as brain, heart, liver, spleen, ovary and testis, in which the MR does not regulate electrolyte homeostasis, evolved early in vertebrate evolution, in a common ancestor of jawed vertebrates. Strong MR expression in ovary and testis suggests a role for Prog-MR complexes in elephant shark reproduction, as well as other novel functions in some other MR-containing tissues. Finally, our data suggest that several 3-ketosteroids, including Prog, may have been ancestral mineralocorticoids.

## Results

### Functional domains of elephant shark MR and other vertebrate MRs

In Figure 1, we compare the functional domains of elephant shark MR to selected vertebrate MRs. Elephant shark MR and human MR have 92% and 67% identity in DBD and LBD, respectively. Interestingly, elephant shark MR has similar conservation to the DBD (91-92%) and LBD (64-69%) of other MRs. The A, B and D domains of elephant shark MR and other MRs are much less conserved.

### Transcriptional activation of full-length and truncated elephant shark MR by corticosteroids, progesterone and spironolactone

We screened a panel of steroids at 0.1 nM and 1 nM for transcriptional activation of full-length and truncated elephant shark MR. At 1 nM, Aldo, cortisol, corticosterone, 11-deoxycorticosterone and 11-deoxycortisol were strong activators of full-length elephant shark MR (Figure 3A) indicating that elephant shark MR has broad specificity for corticosteroids. Interestingly, at 1 nM, 19 norProg has activity comparable to that of the five corticosteroids, while Prog and Spiron have intermediate activity and 17OH-Prog has little activity (Figure 3A).

**Figure 3.**
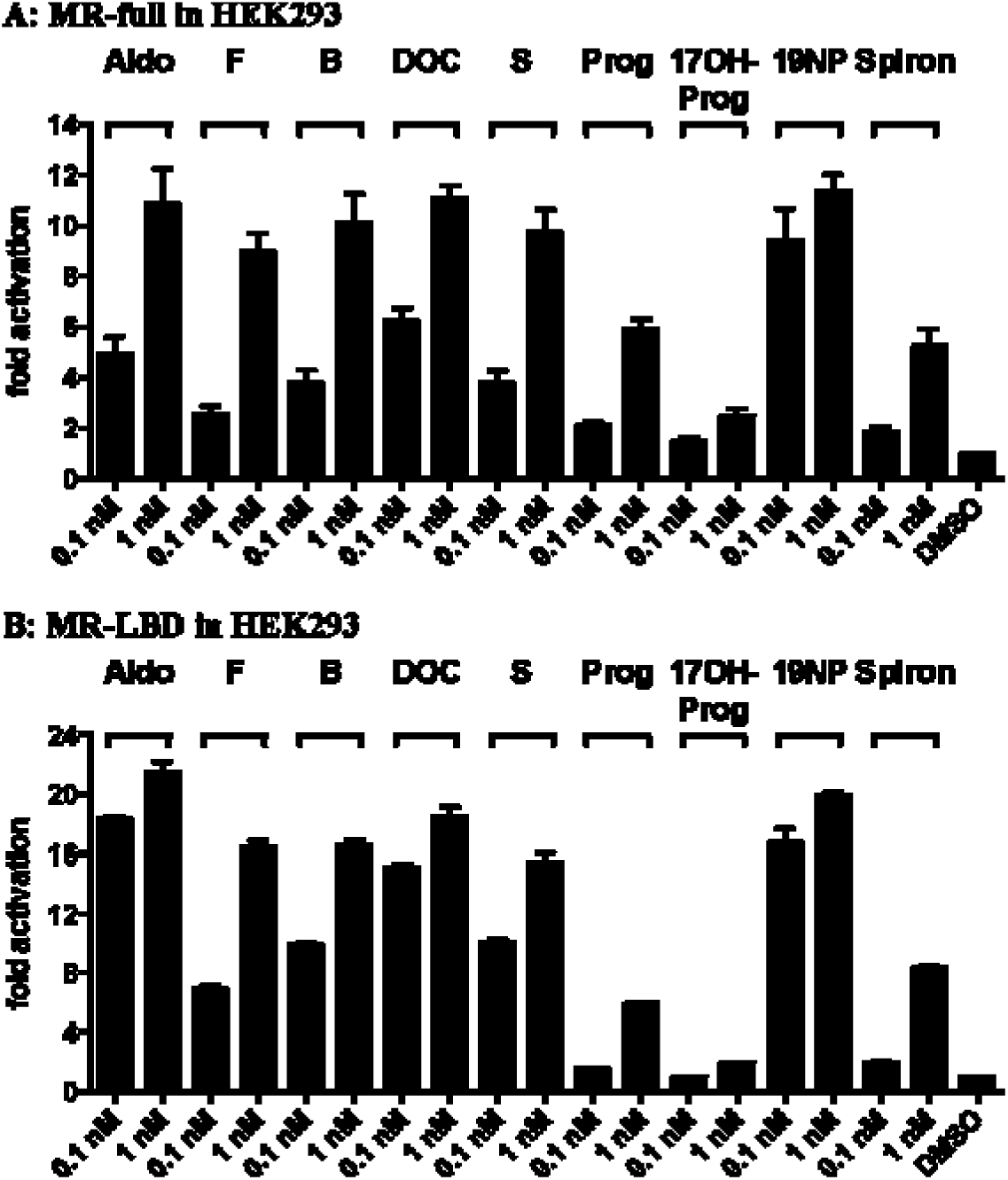
Transcriptional activation of elephant shark MR by 3-ketosteroids. Full length and truncated elephant shark MR were expressed in HEK293 cells with an MMTV-luciferase reporter. **A.** Full length elephant shark MR. Cells were treated with 0.1 nM or 1.0 nM Aldo, cortisol, corticosterone, 11-deoxycorticosterone, 11-deoxycortisol, Prog, 19norProg (19NP), 17OH-Prog, Spiron or vehicle alone (DMSO). **B.** Truncated elephant shark MR. Cells were treated with 0.1 nM or 1.0 nM Aldo, cortisol, corticosterone, 11-deoxycorticosterone, 11-deoxycortisol, Prog, 19norProg (19NP), 17OH-Prog or Spiron or vehicle alone (DMSO). Results are expressed as means ± SEM, n=3. Y-axis indicates fold-activation compared to the activity of control vector with vehicle (DMSO) alone as 1.

In parallel experiments, truncated elephant shark MR, lacking the A/B domain and containing a GAL4-DBD instead of the MR DBD, retained a strong response to all corticosteroids and to 19norProg (Figure 3B). However, Prog and Spiron had reduced activity, while 17OH-Prog had little activity for truncated elephant shark MR.

### EC50 values for steroid activation of elephant shark MR

We determined the concentration-dependence of transcriptional activation of full length elephant shark MR by corticosteroids (Aldo, cortisol, corticosterone, 11-deoxycorticosterone, 11-deoxycortisol) (Figure 4A) and by Prog, 19norProg, 17OH-Prog and Spiron (Figure 4B). We also determined the corresponding concentration-dependent curves for activation of truncated elephant shark MR (Figures 4C and 4D), respectively. Table 1 summarizes the EC50s of corticosteroids for full-length and truncated elephant shark MR. The five corticosteroids, Aldo, corticosterone, 11-deoxycorticosterone, cortisol and 11-deoxycortisol, had similar EC50s and fold activation of full-length elephant shark MR (Figure 4A). As shown in Table 1, EC50s varied from 0.063 nM for 11-deoxycorticosterone to 0.46 nM for cortisol. EC50 values of less than 1 nM are consistent with each corticosteroid being a physiological activator of elephant shark MR. Fold activation compared to Aldo (100%) varied from 83% for 11-deoxycorticosterone and 11-deoxycortisol, to 114% for cortisol (Figure 4A, Table 1), indicating that all five corticosteroids are strong activators of elephant shark MR.

**Table 1.** Corticosteroid activation of full-length MR and GAL4-DBD-MR-LBD.

| MR | Aldo | B | F | DOC | S |
| --- | --- | --- | --- | --- | --- |
|  | EC50 (M) | EC50 (M) | EC50 (M) | EC50 (M) | EC50 (M) |
| **Elephant shark Full** | **1.1 x 10<sup>-10</sup>** | **1.7 x 10<sup>-10</sup>** | **4.6 x 10<sup>-10</sup>** | **6.3 x 10<sup>-11</sup>** | **1.1 x 10<sup>-10</sup>** |
|  | **100%** | **101%** | **114%** | **83%** | **83%** |
| **Elephant shark LBD** | **3.7 x 10<sup>-11</sup>** | **9.9 x 10<sup>-11</sup>** | **1.9 x 10<sup>-10</sup>** | **2.4 x 10<sup>-11</sup>** | **6.8 x 10<sup>-11</sup>** |
|  | **100%** | **90%** | **79%** | **81%** | **77%** |
| <sup>1</sup>**Skate LBD** | **7 x 10<sup>-11</sup>** | **1 x 10<sup>-10</sup>** | **1 x 10<sup>-9</sup>** | **3 x 10<sup>-11</sup>** | **2.2 x 10<sup>-8</sup>** |
| <sup>2</sup>**Human Full** | **2.7 x 10<sup>-10</sup>** | **1.2 x 10<sup>-9</sup>** | **5.5 x 10<sup>-9</sup>** | **4.2 x 10<sup>-10</sup>** | **3.6 x 10<sup>-9</sup>** |
|  | **100%** | **119%** | **133%** | **74%** | **42%** |
| <sup>2</sup>**Human LBD** | **2.8 x 10<sup>-10</sup>** | **5.9 x 10<sup>-10</sup>** | **3. x 10<sup>-9</sup>** | **1.8 x 10<sup>-9</sup>** | **#-** |
|  | **100%** | **95%** | **74%** | **44%** | **\*8%** |
| <sup>2</sup>**Chicken Full** | **6.2 x 10<sup>-11</sup>** | **5.1 x 10<sup>-11</sup>** | **2.8 x 10<sup>-10</sup>** | **3.4 x 10<sup>-11</sup>** | **6.7 x 10<sup>-10</sup>** |
|  | **100%** | **109%** | **128%** | **110%** | **112%** |
| <sup>2</sup>**Chicken LBD** | **1.3 x 10<sup>-10</sup>** | **1.6 x 10<sup>-10</sup>** | **6.9 x 10<sup>-10</sup>** | **1.7 x 10<sup>-10</sup>** | **4.7 x 10<sup>-9</sup>** |
|  | **100%** | **92%** | **75%** | **92%** | **36%** |
| <sup>2</sup>**Alligator-Full** | **2.8x 10<sup>-10</sup>** | **3.6 x 10<sup>-10</sup>** | **6.9x 10<sup>-9</sup>** | **2.3 x 10<sup>-10</sup>** | **2.7 x 10<sup>-9</sup>** |
|  | **100%** | **138%** | **176%** | **85%** | **45%** |
| <sup>2</sup>**Alligator LBD** | **3.5 x 10<sup>-10</sup>** | **3.8 x 10<sup>-10</sup>** | **2.3 x 10<sup>-9</sup>** | **5.2 x 10<sup>-10</sup>** | **#-** |
|  | **100%** | **88%** | **68%** | **51%** | **\*8%** |
| <sup>2</sup>**Xenopus Full** | **4.6 x 10<sup>-10</sup>** | **6.2 x 10<sup>-10</sup>** | **1.1 x 10<sup>-8</sup>** | **7.6 x 10<sup>-10</sup>** | **9.1 x 10<sup>-9</sup>** |
|  | **100%** | **105%** | **126%** | **59%** | **31%** |
| <sup>2</sup>**Xenopus-LBD** | **1.5 x 10<sup>-9</sup>** | **1.9 x 10<sup>-9</sup>** | **1.2 x 10<sup>-8</sup>** | **#-** | **#-** |
|  | **100%** | **74%** | **37%** | **\*10%** | **\*6%** |
| <sup>2</sup>**Zebrafish Full** | **8.2 x 10<sup>-11</sup>** | **3.0 x 10<sup>-10</sup>** | **4.4 x 10<sup>-10</sup>** | **6.3 x 10<sup>-11</sup>** | **4.0 x 10<sup>-10</sup>** |
|  | **100%** | **112%** | **123%** | **103%** | **94%** |
| <sup>2</sup>**Zebrafish LBD** | **2.7 x 10<sup>-11</sup>** | **1.5 x 10<sup>-10</sup>** | **3.1 x 10<sup>-10</sup>** | **1.0 x 10<sup>-10</sup>** | **9.1 x 10<sup>-10</sup>** |
|  | **100%** | **96%** | **77%** | **99%** | **67%** |

<sup>1</sup> (61)<sup>2</sup> (35)

**Figure 4.**
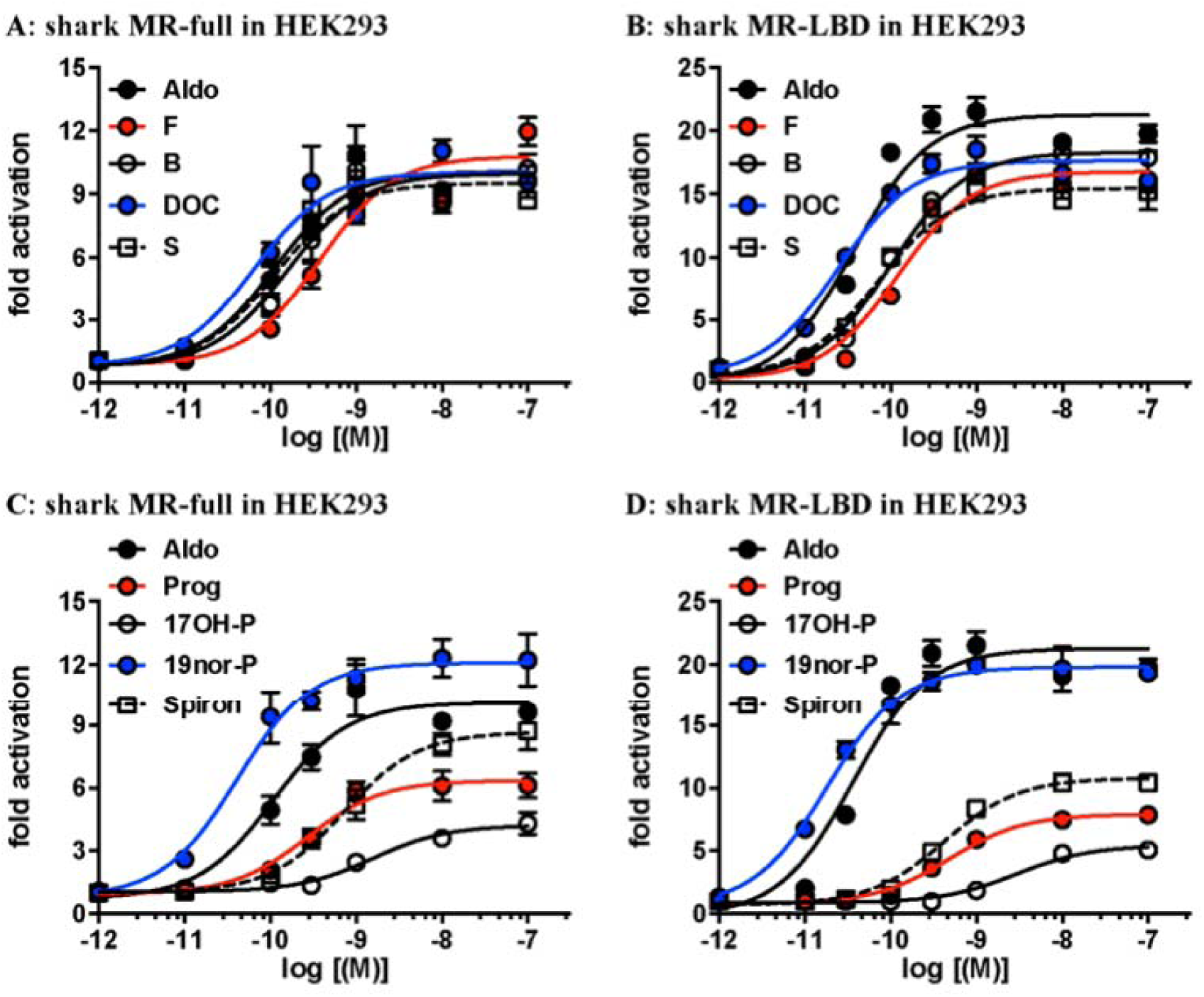
Concentration-dependent transcriptional activation of full-length and truncated elephant shark MR (shark MR) by 3-ketosteroids. Full length and truncated elephant shark MR (shark MR) were expressed in HEK293 cells with an MMTV-luciferase reporter. Full-length elephant shark MR (shark MR) (A) and (C) and truncated elephant shark MR (shark MR) (B) and (D). **(A), (B)** Aldo, cortisol, corticosterone, 11deoxycorticosterone or 11-deoxycortisol. **(C), (D)** Aldo, Prog, 19norProg (19nor-P), 17OH-Prog (17OH-P) or Spiron. Cells transfected with elephant shark MR were treated with increasing concentrations of cortisol, corticosterone, Aldo, 11deoxycorticosterone, 11-deoxycortisol, Prog, 19norProg, 17OH-Prog, Spiron or vehicle alone (DMSO). Y-axis indicates fold-activation compared to the activity of control vector with vehicle (DMSO) alone as 1.

Truncated elephant shark MR also had a strong response to corticosteroids, which had EC50 values from 0.024 nM for 11-deoxycorticosterone to 0.19 nM for cortisol (Table 1). Fold activation compared to that for Aldo (100%), decreased to 79% for cortisol, 81% for 11-deoxycorticosterone and 77% for 11-deoxycortisol.

Table 1 also contains, for comparison, previously determined EC50s of corticosteroids for full-length and truncated human, chicken, alligator, Xenopus and zebrafish MRs (*35*) and skate MR (*61*). Unlike elephant shark MR, there are differences among terrestrial vertebrate MRs in their EC50s and fold activation for the five glucocorticoids. Aldo and corticosterone are good activators of full-length and truncated terrestrial vertebrate MRs, while 11-deoxycorticosterone and 11-deoxycortisol are weak activators of truncated human, alligator and *Xenopus* MRs (*35*). Interestingly, all corticosteroids have low EC50s and strong fold activation for full-length and truncated zebrafish MR (Table 1) (*35*) indicating that zebrafish MR retains responses to corticosteroids found in elephant shark MR.

Among the progestins, we find that 19norProg is the most active for full-length and truncated elephant shark MR (Figure 4D, Table 2). 19norProg has EC50s of 0.43 nM for full-length and 0.018 nM truncated for elephant shark MR. This is comparable to the EC50 values of corticosteroids for full-length and truncated elephant shark MR. Fold activation, of 19norProg for full-length and truncated elephant shark MR was 84% and 98% respectively compared to that for Aldo (100%), indicating that 19norProg could be a physiological activator of elephant shark MR. Prog and Spiron have strong EC50s of 0.27 nM and 0.66 nM, respectively, for full-length elephant shark MR. However, activation decreased to about 45% of Aldo’s fold activation of full-length MR. 17OH-Prog had an EC50 of 1.9 nM and only 25% of Aldo’s fold activation of full-length MR.

**Table 2.**
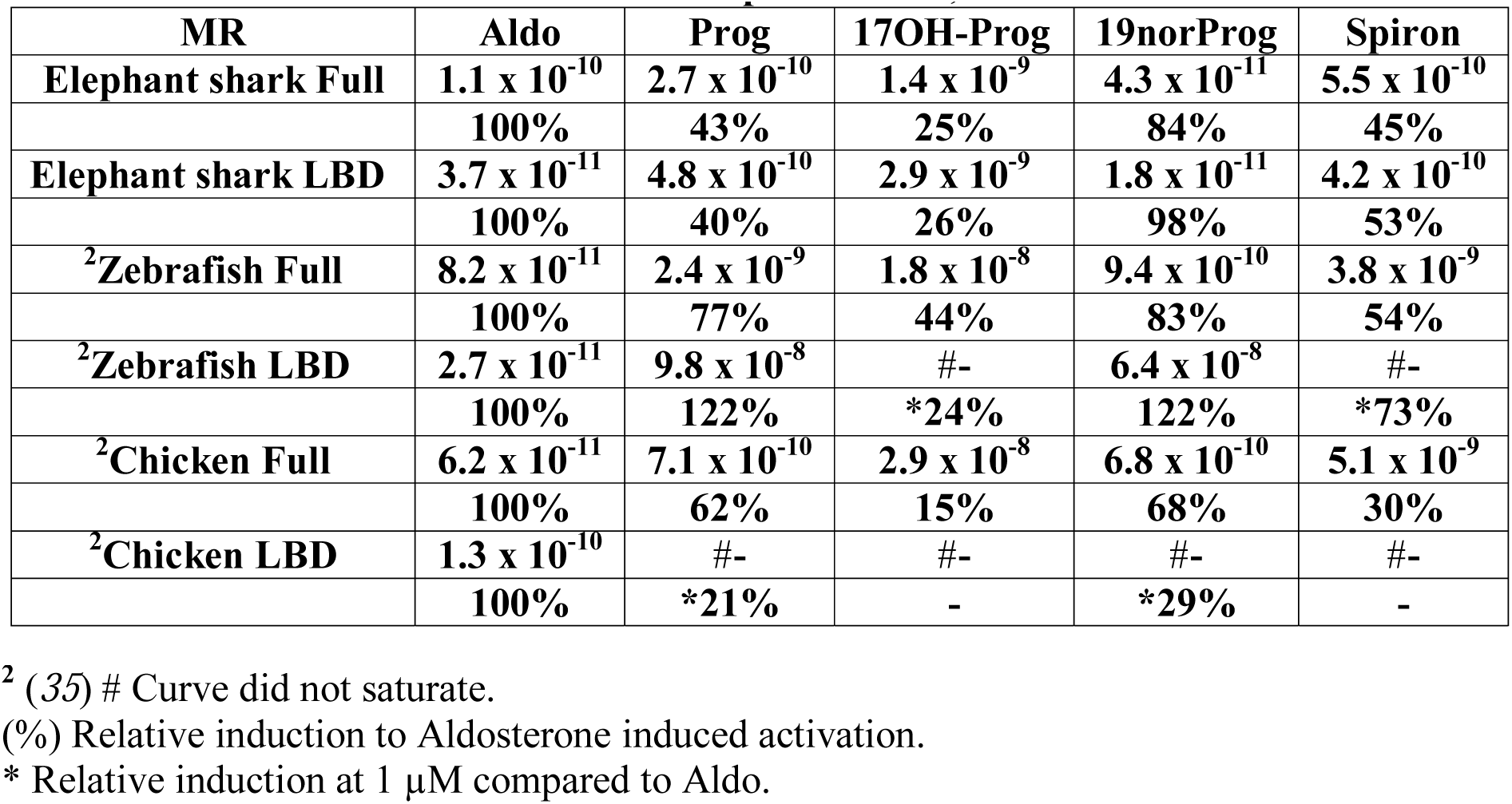
EC50 values for progestin and spironolactone activation of full-length and GAL4-DBD-MR-LBD constructs of elephant shark, zebrafish and chicken MR.

| MR | Aldo | Prog | 17OH-Prog | 19norProg | Spiron |
| --- | --- | --- | --- | --- | --- |
| Elephant shark Full | $1.1 \times 10^{-10}$ | $2.7 \times 10^{-10}$ | $1.4 \times 10^{-9}$ | $4.3 \times 10^{-11}$ | $5.5 \times 10^{-10}$ |
|  | 100% | 43% | 25% | 84% | 45% |
| Elephant shark LBD | $3.7 \times 10^{-11}$ | $4.8 \times 10^{-10}$ | $2.9 \times 10^{-9}$ | $1.8 \times 10^{-11}$ | $4.2 \times 10^{-10}$ |
|  | 100% | 40% | 26% | 98% | 53% |
| <sup>2</sup>Zebrafish Full | $8.2 \times 10^{-11}$ | $2.4 \times 10^{-9}$ | $1.8 \times 10^{-8}$ | $9.4 \times 10^{-10}$ | $3.8 \times 10^{-9}$ |
|  | 100% | 77% | 44% | 83% | 54% |
| <sup>2</sup>Zebrafish LBD | $2.7 \times 10^{-11}$ | $9.8 \times 10^{-8}$ | #- | $6.4 \times 10^{-8}$ | #- |
|  | 100% | 122% | *24% | 122% | *73% |
| <sup>2</sup>Chicken Full | $6.2 \times 10^{-11}$ | $7.1 \times 10^{-10}$ | $2.9 \times 10^{-8}$ | $6.8 \times 10^{-10}$ | $5.1 \times 10^{-9}$ |
|  | 100% | 62% | 15% | 68% | 30% |
| <sup>2</sup>Chicken LBD | $1.3 \times 10^{-10}$ | #- | #- | #- | #- |
|  | 100% | *21% | - | *29% | - |

<sup>2</sup> (35) # Curve did not saturate.
(%) Relative induction to Aldosterone induced activation.
\* Relative induction at 1 $\mu$ M compared to Aldo.

Previously we reported that Prog, 19norProg and Spiron are transcriptional activators of full-length and truncated chicken and zebrafish MRs (*35*). However, EC50 values of Prog, 19norProg and Spiron for these MRs were not determined. We have remedied this omission and report their EC50 values, as well as EC50 values for 17OH-Prog, in Table 2 and Figure 5, for comparison with full-length and truncated elephant shark MR.

**Figure 5.**
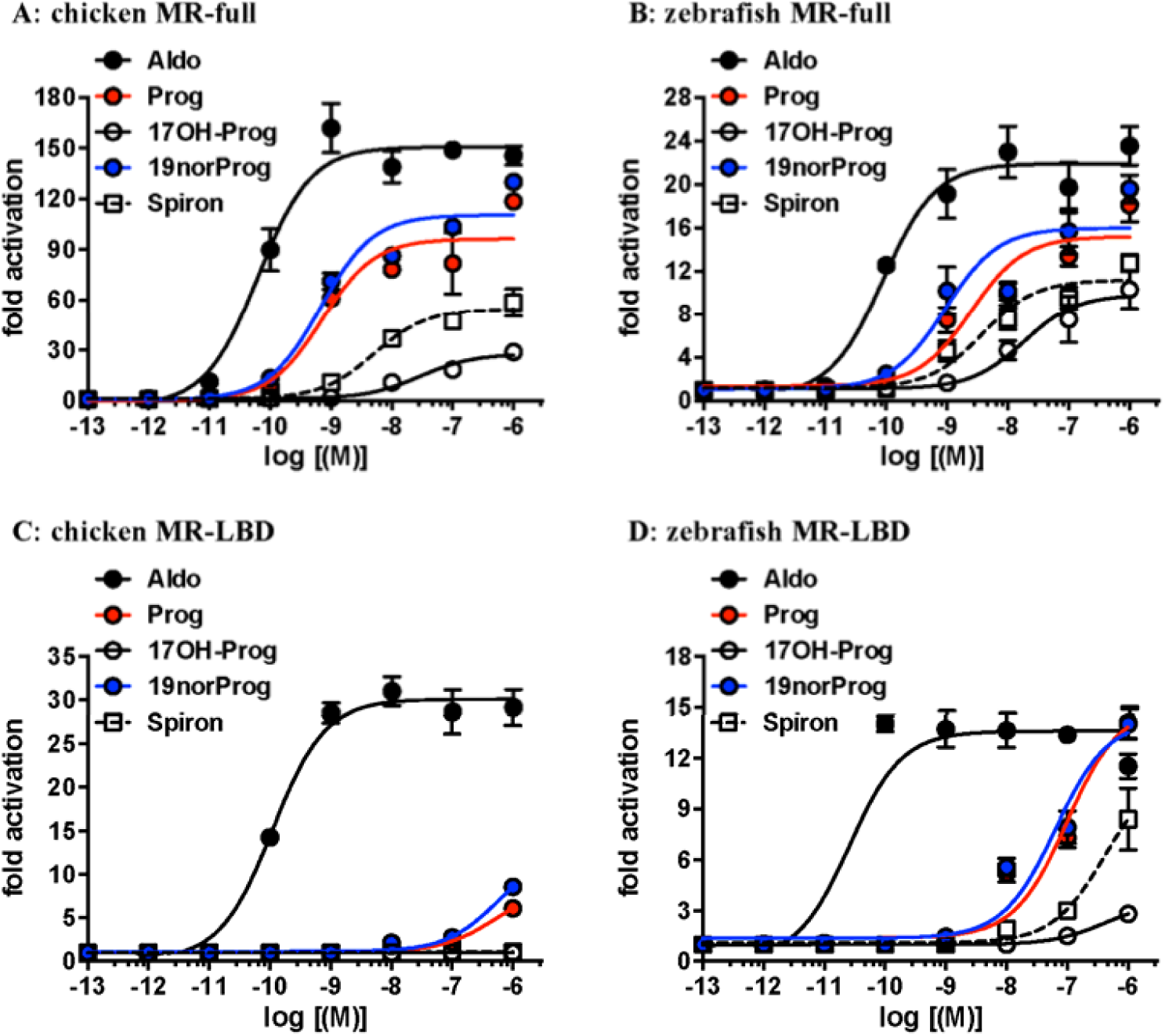
Concentration-dependent transcriptional activation of full-length and truncated chicken and zebrafish MR by progestins and Spironolactone. HEK293 cells transfected with chicken and zebrafish MR were treated with increasing concentrations of Prog, 19norProg, 17OH-Prog or Spiron. Full-length chicken MR (A) and zebrafish MR (B) and truncated chicken MR (C) and zebrafish MR (D). Y-axis indicates fold-activation compared to the activity of control vector with vehicle (DMSO) alone as 1.

Regarding full-length chicken MR, Prog and 19norProg have EC values of 0.68 nM and 0.71 nM, respectively, which are low enough for physiological activation of chicken MR (Figure 5A, Table 2). Fold activation by Prog and 19norProg is 62% and 68%, respectively, of Aldo (100%). 17OH-Prog has an EC50 of 29 nM and fold-activation that is 15% of Aldo (100%). Spiron has an EC50 of 5.1 nM for full-length chicken MR. Activation of truncated chicken MR by Prog,19norProg, 17OH-Prog and Spiron (Figure 5C, Table 2) was too low for calculation of EC50 values, suggesting that allosteric interactions between the NTD and LBD are important in progestin activation of chicken MR.

Regarding zebrafish MR, 19norProg, had an EC50 of 0.9 nM and 83% of Aldo’s fold activation, and Prog had an EC50 of 2.4 nM and 77% of Aldo’s fold activation for full-length zebrafish MR. These responses are sufficient for both 19norProg and prog to be physiological activators of zebrafish MR. In contrast, 17OH-Prog had an EC50 of 18 nM and fold activation that 44% of Aldo’s activation of full-length zebrafish MR. EC50s for truncated zebrafish MR were higher for all progestins and Spiron. The EC50s of Prog and 19norProg were 98 nM and 64 nM respectively, which is much weaker than for full-length zebrafish MR. Activation of truncated zebrafish MR by 17OH-Prog and Spiron was too weak for their EC50s of to be calculated. This contrasts with strong activation by corticosteroids of full-length zebrafish MR (Table 1), (Figure 5, Table 2), suggesting that allosteric interactions between the NTD/DBD and LBD contribute to progestin activation of zebrafish MR.

### RNA-seq analysis of elephant shark MR

We examined expression of level of elephant shark MR gene in 10 tissues based on RNA-seq data (Figure 6A). The MR gene was found to express widely in all tissues, including gills and kidney, two traditional mineralocorticoid-responsive tissues. Interestingly there was considerably higher expression in the ovary and testis, the two reproductive tissues analyzed.

**Figure 6A.**
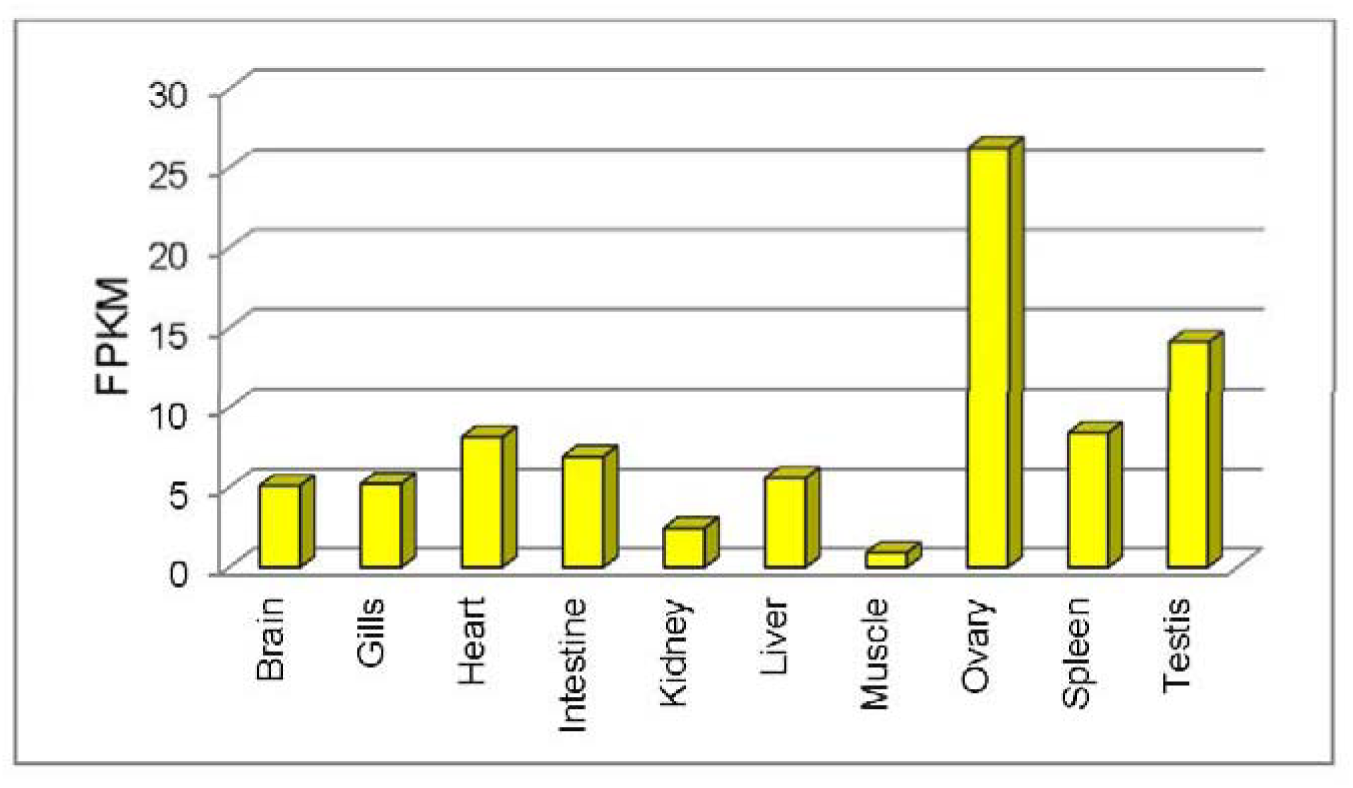
Expression level of MR gene in 10 tissues of elephant shark estimated based on RNA-seq data. Transcript abundances are shown in terms of normalized counts called Fragments per kilobase of exon per million fragments mapped (FPKM) (*62*). FPKM is estimated by normalizing gene length followed by normalizing for sequencing depth.

### RNA-seq analysis of human MR

RNA-seq analysis of human MR (*63*) (Figure 6B) reveals that the MR is expressed in kidney, colon, brain, heart, liver, ovary, spleen and testis. This pattern of expression of human MR in diverse tissues is similar to that of elephant shark MR.

**Figure 6B.**
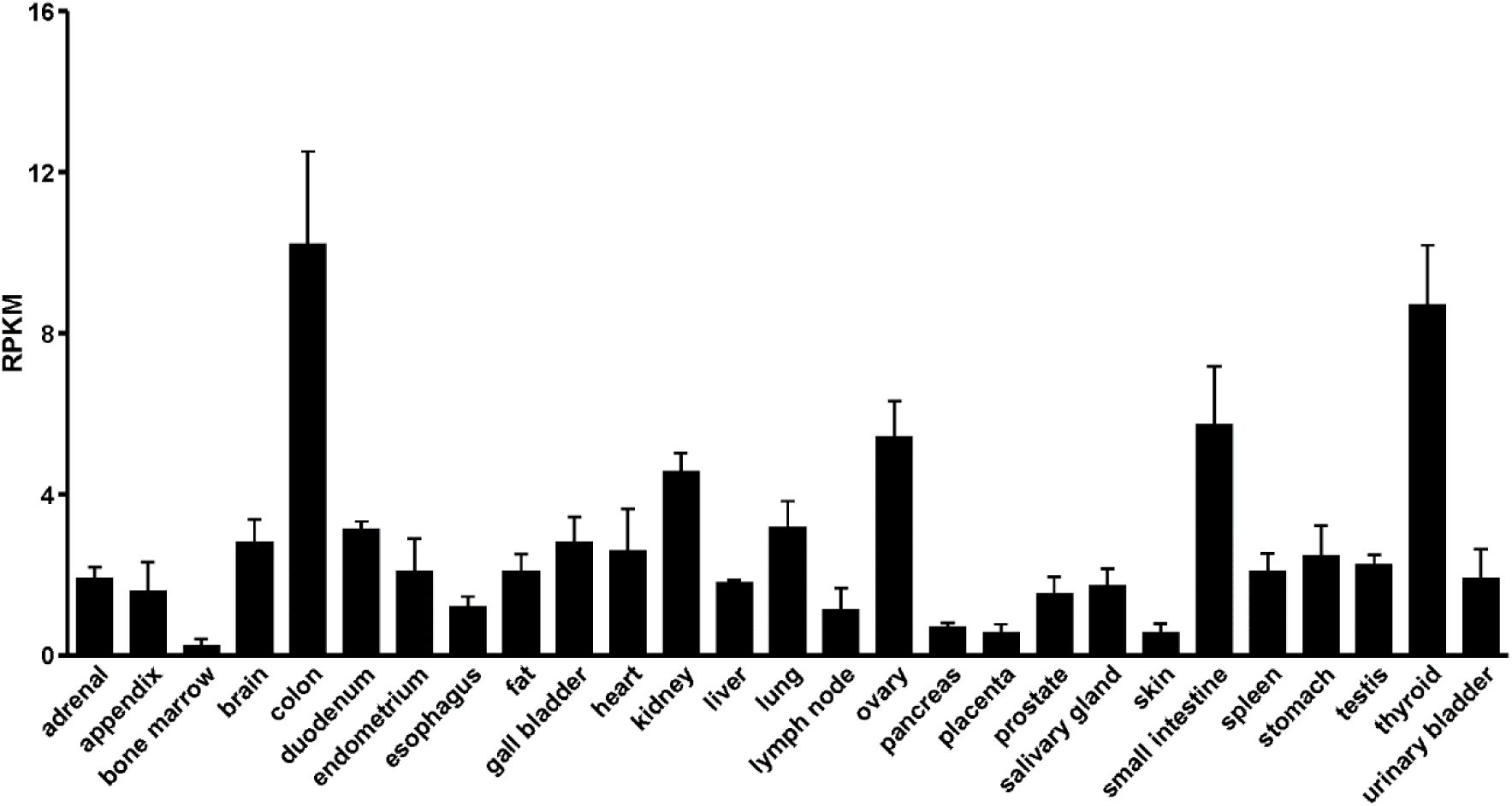
Expression level of human MR based on RNA-seq data. Transcript abundances are shown in terms of normalized counts called Reads Per Kilobase of transcript per Million mapped reads (RPKM) (*63*).

## Discussion

Cartilaginous fish, including elephant sharks, occupy a key position in the evolution of vertebrates as an out-group to ray-finned fish, the largest group of extant vertebrates, and the lobe-finned fish, which are the forerunners of terrestrial vertebrates. Importantly, the elephant shark genome is evolving slowly (*9*), making it attractive for studying ancestral proteins, including the MR, which first appears as a distinct MR ortholog in cartilaginous fish (*5, 6, 10, 61*).

Our investigation of corticosteroid activation of elephant shark MR reveals that Aldo, corticosterone, 11-deoxycorticosterone, cortisol and 11-deoxycortisol have EC50s below 1 nM for full-length elephant shark MR, (Figure 4, Table 1). Prog, 19norProg and Spiron also have sub-nM EC50s for full-length elephant shark MR. In addition to their low EC50s, all of these corticosteroids and 19norProg have a similar strong fold activation of transcription of full-length MR (Figure 4A, C), while Prog has about 43% of fold activation by Aldo (100%). Thus, several corticosteroids, as well as 19norProg and Prog are potential physiological mineralocorticoids for elephant shark MR.

Compared to full-length elephant shark MR, the EC50s of all five corticosteroids and 19norProg are lower for its truncated MR, while the EC50 for Spiron is slightly lower, and the EC50s for Prog and 17OH-Prog are about 2-fold higher (Table 1, Table 2).

Regarding truncated skate MR, most of the EC50s of corticosteroids (*61*) are similar to that for elephant shark MR (Table 1). The exception is 11-deoxycortisol, which has over 200-fold higher EC50 for skate MR than for elephant shark MR.

### Progesterone and/or 19norProgesterone may be mineralocorticoids in cartilaginous fishes

The Prog concentration in female elephant sharks is 4.4 ng/ml (14 nM) (*64*). In draughtboard sharks (*Cephaloscyllium laticeps*), Prog levels in females are 8 ng/mL (25.4 nM) and in males are 1 ng/ml (3.2 nM) (*65*). In female zebrasharks (*Stegostoma fasciatum*), Prog levels are 10 ng/ml (31.8 nM) (*66*). Together this indicates that Prog concentrations are sufficient to activate the MR in cartilaginous fishes.

19norProg has an EC50 of 0.043 nM for elephant shark MR. Moreover, 19norProg evokes a stronger response from elephant shark MR than Aldo (Figure 4A, B). C19 demethylase activity has been found in mammalian kidney (*59*). If C19 demethylase is present in elephant shark, then 19norProg needs to be considered as a potential physiological mineralocorticoid.

Although in cell assays 19norProg is an agonist for elephant shark, zebrafish and chicken MRs (Table 2), at 1 nM, 19norProg is an antagonist for human MR (*35, 47*). This contrasts with *in vivo* studies in rats, which found that 19norProg is an MR agonist with about 100-fold weaker activity than Aldo (*59, 60*) A possible explanation for this difference is that in rats, 19norProg is metabolized to 19nor-11-deoxycorticosterone, which is a mineralocorticoid agonist (*67–69*). Another possibility, based on 11β-hydroxyprogesterone activation of human MR (*27*), is that 19norProg is metabolized to 11β-hydroxy19norProg.

We propose that transcriptional activation of elephant shark MR by 19norProg, as well as by Prog and Spiron, can be explained by Geller et al.’s (*47*) discovery that S810L human MR mutant is activated by 1 nM Prog, 19norProg, and Spiron, unlike wild-type human MR, in which these steroids are MR antagonists. Based on a 3D model of S810L MR Geller et al. (*47*) proposed that a contact between Leu-810 and Ala-773 was sufficient for transcriptional activation of S810L MR by Prog and 19norProg. This motivated Geller et al. (*47*) to construct a human S810M mutant, which was activated by 19norProg. Elephant shark MR and skate MR contain a methionine at this position corresponding to Ser810 and an alanine corresponding to Ala-773 (Figure 7). Based on Geller et al.’s model, we propose that transcriptional activation of elephant shark MR by 19norProg is due to a contact between Met-782 (helix 5) and Ala-745 (helix 3), which stabilizes the A ring of 19norProg, promoting transcriptional activation.

**Figure 7.**
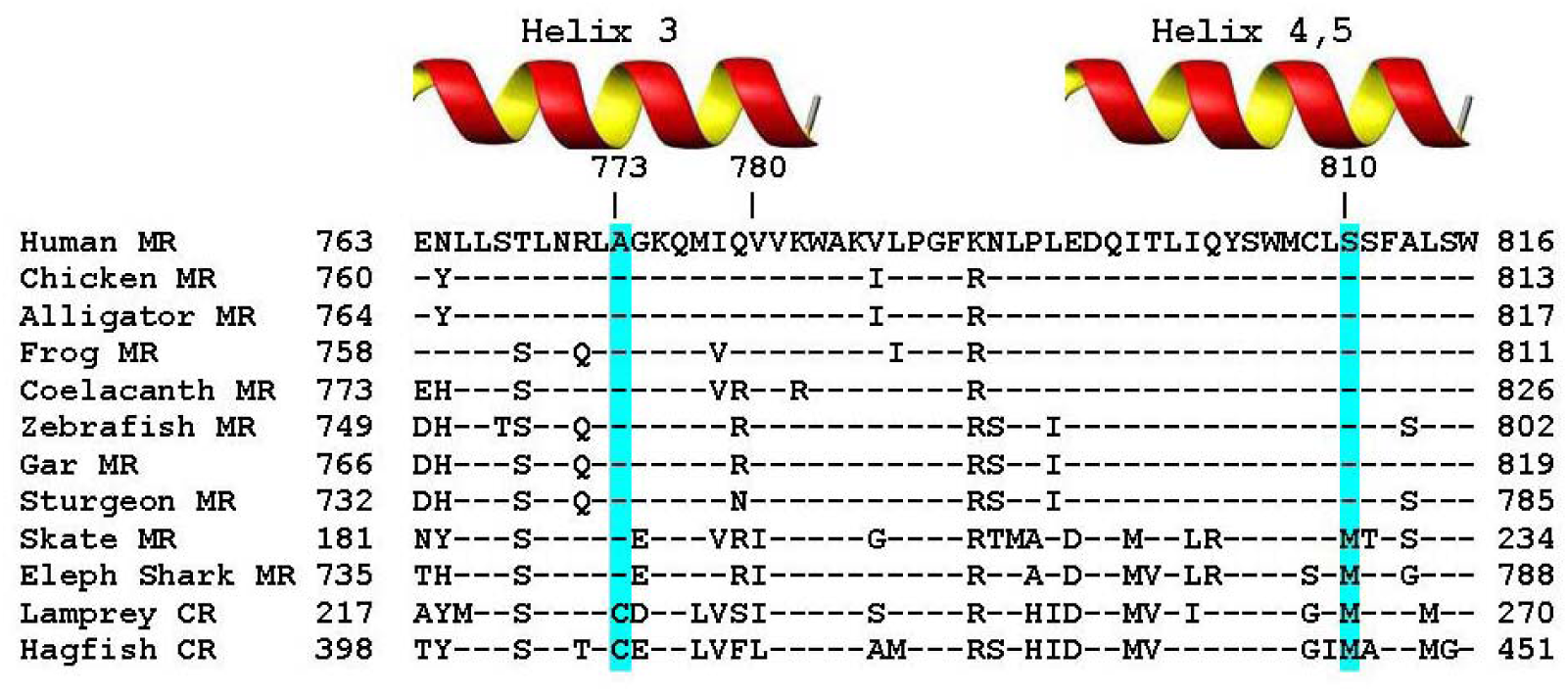
Alignment of elephant shark MR to Serine-810 and Alanine-773 in helices 3-5 in human MR. Elephant shark MR and skate MR contain a methionine corresponding to human Ser-810 and an alanine corresponding to Ala-773. Lamprey CR and hagfish CR also contain a corresponding methionine, as well as a cysteine corresponding to Ala-773. Human Ser-810 and Ala-773 are conserved in coelacanths, terrestrial vertebrate and ray-finned fish MRs. Amino acids that are identical to amino acids in human MR are denoted by (-).

### Role for elephant shark MR in reproductive physiology

Prog’s EC50 of 0.27 nM for elephant shark MR, Prog’s physiological concentration of 14.5 nM (50) and high MR expression in elephant shark ovary and testis (Figure 6A) suggests that a Prog-MR complex is important in reproductive responses in elephant shark. Of course, Prog also acts as a reproductive steroid in ovary and testis via transcriptonal activation of the PR (58, 59). Based on evidence that Prog activates the MR in several ray-finned fishes (24, 36, 44, 46), a Prog-MR complex also may be active in reproductive tissues and other tissues in ray-finned fish, as well as in cartilaginous fish.

RNA-seq analysis also finds MR expression in elephant shark gills and kidneys (Figure 6A), two classical targets for MR regulation of electrolyte transport (6). Moreover, RNA-seq analysis also identifies MR expression in elephant shark heart and brain, two other tissues in which corticosteroids have important physiological actions via the MR (16-18, 20, 21, 60-62). Indeed, RNA seq analysis of elephant shark MR indicates that expression the MR in diverse tissues was conserved during the descent from cartilaginous fishes to humans. Expression of shark MR in many tissues (brain, heart, liver, ovary) in which the MR is not likely to regulate electrolyte homeostasis, the MR’s classical function, further supports evidence from the last thirty years (3, 17-19, 60, 62-64) that mineralocorticoid activity is an incomplete functional description of this nuclear receptor. An alternative name is needed to describe more completely the functions of the MR.

## Materials and Methods

### Chemical reagents

Aldo, cortisol, corticosterone, 11-deoxycorticosterone, 11-deoxcortisol, Prog, 19norProg, 17OH-Prog and Spiron were purchased from Sigma-Aldrich. For reporter gene assays, all hormones were dissolved in dimethylsulfoxide (DMSO); the final DMSO concentration in the culture medium did not exceed 0.1%.

### Construction of plasmid vectors

The full-coding regions from elephant shark MR were amplified by PCR with KOD DNA polymerase. The PCR products were gel-purified and ligated into pcDNA3.1 vector (Invitrogen) for the full-coding region or pBIND vector for D-E domains (*70*).

### Elephant shark MR gene expression analysis

We had previously generated RNA-seq for several tissues of elephant shark as part of the elephant shark genome project (*9*) and submitted them to NCBI (accession number SRA054255). We downloaded RNA-seq reads for brain, gills, heart, intestine, kidney, liver, muscle, ovary, spleen, and testis, and assembled each of them into transcripts using the program Trinity, version r2013-08-14 (*71*). The assembled transcripts were used to determine the expression level of MR genes as detailed below. To determine the expression level of MR genes, we performed abundance estimation of transcripts from the afore mentioned 10 tissues. Trinity transcripts from all ten tissues and full-length cDNA sequence of the MR gene were combined together and clustered using CD-HITv4.6.1 at 100% identity (*72*). RNA-seq reads from each of the ten tissues was independently aligned to the clustered transcript sequences and abundance of MR gene transcripts was estimated by RSEMv1.2.25 (*62*) which uses bowtiev2.2.6 for aligning (*73*). Transcript abundances were measured in terms of normalized counts called Fragments per kilobase of exon per million fragments mapped (FPKM) (*62*). FPKM is estimated by normalizing the gene length followed by normalizing for sequencing depth.

### Transactivation Assay and Statistical Methods

Transfection and reporter assays were carried out in HEK293 cells, as described previously (*70, 74*). All experiments were performed in triplicate. The values shown are mean ± SEM from three separate experiments, and dose-response data and EC50 were analyzed using GraphPad Prism. Comparisons between two groups were performed using *t*-test, and all multi-group comparisons were performed using one-way ANOVA followed by Bonferroni test. *P* < 0.05 was considered statistically significant. The use of Hek293 cells and an assay temperature of 37 C does not replicate the physiological environment of Elephant sharks. Nevertheless, studies with Hek293 cells and other mammalian cell lines have proven useful for other studies of transcriptional activation by corticosteroids of skate (*61*) and teleost fish (*36, 44, 75, 76*) MRs.

## Acknowledgments

**Funding**

K.O. was supported by the Japan Society for the Promotion of Science (JSPS) Research Fellowships for Young Scientists. This work was supported in part by Grants-in-Aid for Scientific Research 23570067 and 26440159 (YK) from the Ministry of Education, Culture, Sports, Science and Technology of Japan. M.E.B. was supported by Research fund #3096. BV was supported by the Biomedical Research Council of A*STAR, Singapore.

We thank Dr. Cynthia Awruch for advice on progesterone levels in cartilaginous fishes.

## Author contributions

Y.K., S.K., K.O., X.L., S.O. and N.E.P. carried out the research. Y.K. and S.K. analyzed data. W.T. and S.H. aided in the collection of animals. NEP assembled and analyzed RNAseq data. Y.K. and M.E.B. conceived and designed the experiments. Y.K., M.E.B. and B.V. wrote the paper. All authors gave final approval for publication.

